# Population regulation in microbial consortia using dual feedback control

**DOI:** 10.1101/120253

**Authors:** Xinying Ren, Ania-Ariadna Baetica, Anandh Swaminathan, Richard M. Murray

**Affiliations:** Department of Control and Dynamical Systems, California Institute of Technology, Pasadena, CA..; Department of Biology and Biological Engineering, California Institute of Technology, Pasadena, CA.

## Abstract

An ongoing area of study in synthetic biology has been the design and construction of synthetic circuits that maintain homeostasis at the population level. Here, we are interested in designing a synthetic control circuit that regulates the total cell population and the relative ratio between cell strains in a culture containing two different cell strains. We have developed a dual feedback control strategy that uses two separate control loops to achieve the two functions respectively. By combining both of these control loops, we have created a population regulation circuit where both the total population size and relative cell type ratio can be set by reference signals. The dynamics of the regulation circuit show robustness and adaptation to perturbations in cell growth rate and changes in cell numbers. The control architecture is general and could apply to any organism for which synthetic biology tools for quorum sensing, comparison between outputs, and growth control are available.

## I. INTRODUCTION

A primary area of study in synthetic biology has been the implementation of synthetic gene circuits with novel functionality in single cells. The first synthetic gene circuits included oscillators [1], [2] and toggle switches [3], [4]. In oscillator circuits, the expression of a gene of interest oscillates repeatedly over time. In toggle switch circuits, a gene’s expression can be switched between two stable steady state levels. In both cases, the circuit is implemented at the single cell level. Recent applications include the engineering of metabolic pathways in single cells to produce fuels [5] or drugs [6] and the manipulation of a cell’s DNA to implement state machines in single cells [7], [8].

However, there are challenges associated with implementing biological circuits in single cells. One challenge is that when composing genetic circuit parts into larger circuits, loading effects from a downstream module can negatively impact the performance of an upstream module. This phenomenon, termed retroactivity, can interfere with circuit behavior when attempting to build complex synthetic gene circuits [9]. Moreover, single cells have limited pools of resources. Complex circuits with many parts use more cellular resources, and hidden interactions that arise through resource competition can also negatively impact circuit performance [10], [11].

Implementing genetic circuits in multiple cells alleviates these two challenges. With different circuit components in different cells, the components have separate resource pools and thus cannot compete for resources. In addition, the communication between cells is solely mediated by small molecules that typically exist at very large copy numbers, which mitigates loading effects from retroactivity. This approach is illustrated in [12], where the authors use different combinations of yeast strains to implement different logical functions.

Previously developed synthetic gene circuits that function at the population level and involve feedback include a population control circuit that regulates the number of cells in a culture [13], a predator prey system with two cell strains [14], a two strain system for programmed pattern formation [15], and a two strain population level oscillator [16]. However, there are challenges associated with implementing circuits at the population level across multiple cell strains. One of these challenges is maintaining a stable population fraction of all cell strains. When implementing a two strain system, the ratio between the two cell types might require tuning for the best performance. In addition, there might also be an optimal total population size as too many cells would deplete the resources of the consortium.

Here, we present a control strategy for tuning the cell type ratio as well as the total population size in a two strain system. By using two separate control loops to control the total cell number and the ratio between the two cell types, we demonstrate that both the total population size and the cell type ratio are independently tunable. We show that our control architecture implements a lag compensator. Furthermore, we show that the total population size and cell type ratio are robust to perturbations in the number of cells of either strain and are also robust to perturbations in the growth rate of either strain.

The organization of the paper is as follows. In Section 2, we give an overview of the biological background for the problem and we introduce our design strategy of using two separate loops to control the total population size and the cell type ratio. In Section 3, we provide a model of the control loop for maintaining the total population size, and we demonstrate its effectiveness. We also show that the global population size control loop implements a lag compensator. In Section 4, we introduce a model for the control loop that maintains cell type ratio and again show that the controller implements a lag compensator. We demonstrate that the cell type ratio is robust to perturbations in cell growth rate. In Section 5, we combine the two loops into one model and show that total population size and cell type ratio are independently tunable. We summarize the main findings of the paper and discuss future work in the Discussion section.

## II. THE DESIGN STRATEGY

Our proposed population regulation circuit in microbial consortia consists of two feedback control loops. The global regulation and the co-regulation both involve a controller regulating either cell growth or death processes. By coupling the two loops, we can achieve separate functions that simultaneously regulate the absolute population count and the relative ratio between the two cell strains.

### A. Biological background

In order to control growth of different cells strains, we require biological sensors, comparators, and actuators [17]. The sensors need to sense the population size, the comparators need to compare the population size to a reference signal, and the actuators need to use the output from the comparators to drive cell growth such that the error between the reference population size and the actual population size is reduced. Here, we briefly describe synthetic biological systems that can implement each of these three crucial functions.

Quorum sensing systems in bacteria can be used as sensors for population size. In quorum sensing systems, each cell constitutively produces and secretes a small signal molecule, so the concentration of signaling molecules in solution is proportional to the population size [18]. Downstream gene expression machinery responds to the concentration of the signaling molecules in a graded fashion. While quorum sensing systems are most commonly used in bacteria, similar tools exist in yeast [12] and in mammalian systems [19].

To compare the sensed population size to the reference population size, we need gene circuits that can subtract the two quantities in a chemical manner. This can be achieved by using two proteins, where one protein sequesters the other and inhibits its function. This type of system can be constructed using engineered protein scaffolds [20] or it can be leveraged from a natural system that already exists [21]. Systems that inhibit gene expression at the RNA level can also provide similar functionality [22].

Finally, the difference between the measured and the reference signals must be used to actuate cell growth in order to modulate the population size. Typically, cell growth actuation strategies depend on modulating the expression of a gene that is essential for cell growth. When expression of the essential gene is decreased, the cells grow more slowly. The gp2 phage protein stops bacterial growth by inhibiting bacterial RNA polymerase [23]. Similarly, using an inducible RNA polymerase allows control of cell growth by controlling RNA polymerase expression [24]. Another method that allows for cell growth control is toxin-antitoxin systems. In these systems, a toxin protein slows down cell growth or kills the cell, while an antitoxin protein sequesters the toxin and inhibits its toxicity [21]. Toxin-antitoxin systems are especially useful for building growth controllers, as they can be employed as comparators and actuators.

In this paper, we present a general control design that should be applicable to any synthetic biology organism where the appropriate tools for sensing, comparison, and actuation are available. However, our specific inspiration is to achieve growth control in *E. coli* using quorum sensing [18] for sensing, the ccdB/ccdA toxin-antitoxin system [21] and RNA antisense technology [22] for comparison, and the ccdB toxin and the gp2 protein [23] for actuation.

### B. The global regulation loop for total population control

The global regulation systemcontrols the total population of all strains in the culture and consists of three modules. The cell dynamics module includes the growth and division processes of the cell. The communication module relies on a global quorum sensing system where all cell strains produce and sense a common signal molecule. The feedback controller module is designed to ensure homeostasis of the total cell population by comparing the output and the reference and by actuating the corresponding cell growth process to decrease the error.

The biological design of the global regulation loop is illustrated in Figure 1a. The reference is set by the induction rate of biochemical species *G*, which activates the cell growth and division processes. Species *G* can be strongly sequestered and inactivated by species *D* to decrease cell growth rate. All cell strains release and sense a common signal molecule *S*_*g*_ in a global quorum sensing system. When the total population of all cells increases, more signal molecules *S_g_* are synthesized and released into the environment. These signals diffuse across membranes into cells and activate reactions that produce species *D*. Therefore, more species *D* molecules bind with species *G* molecules and inhibit the cell growth. This negative feedback enables the total cell population to maintain a steady state that tracks the reference signal, which can be set by tuning the basal induction of *G*.

**Fig. 1.**
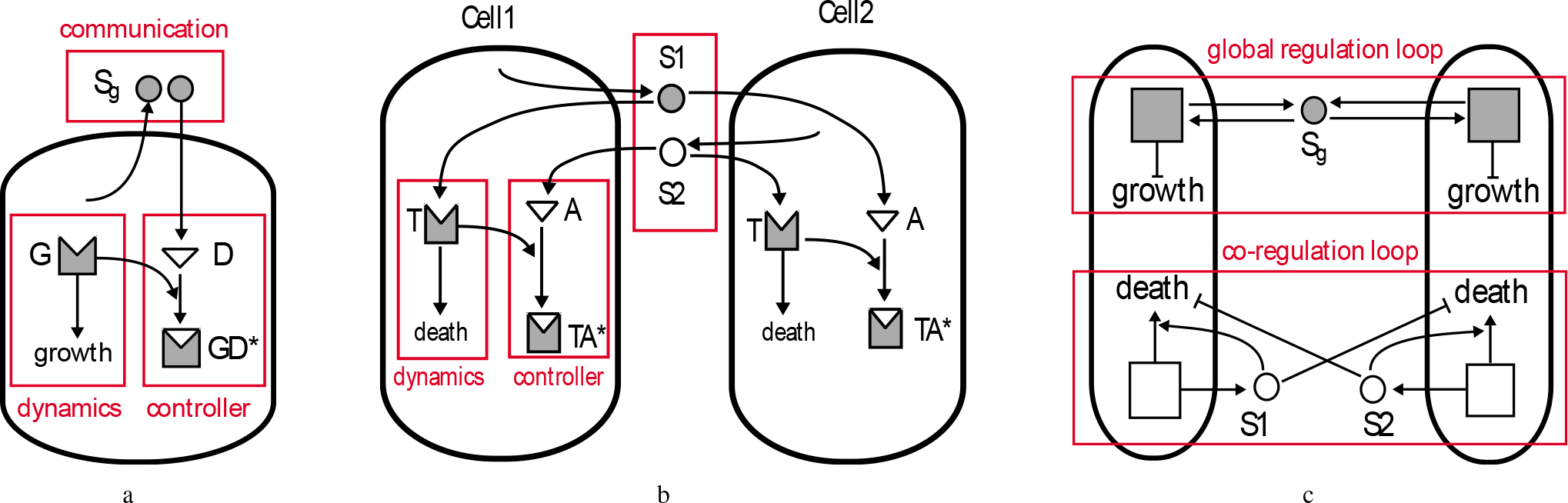
Biological design of the global regulation loop, of the co-regulation loop, and of the dual loop. a. It describes the global regulation that controls the total population. The reference signal is set by internal tuning of the induction rate of species *G*. Cells release and sense a global quorum sensing signal molecule *S_g_* and the concentration of *S_g_* depends on the total population. Signal *S*_*g*_ diffuses inside cells and it activates the production of species *D*, which binds to species *G* to inhibit cell growth and division. b. It describes the co-regulation that controls the relative population ratio between two cell strains. Cell strain Cell_1_ releases and senses a quorum sensing signal molecule *S*_1_ that diffuses and activates the production of toxin *T* in Cell_1_. Cell_1_ also senses another orthogonal signal molecule *S*_2_ released by cell strain Cell_2_ to activate the production of antitoxin A, which binds with *T* to inhibit the killing process in Cell_1_. The negative feedback controller in Cell_1_ regulates the death dynamics to track the population of Cell_2_ and the similar design in Cell_2_ tracks population of Cell_1_ in turn. c. It describes the dual loop regulation that couples both global and co-regulation to realize the function of simultaneously controlling the total population and the relative ratio of the two cell strains.

### C. The co-regulation loop for relative population ratio control

To regulate the relative population ratio between two cell strains, we design a co-regulation loop consisting of a cell dynamics module that regulates cell death, a communication module of two orthogonal quorum sensing systems, and a feedback controller module, which compares the difference between populations of two strains and actuates the antitoxin production in the feedback.

As illustrated in Figure 1b, we consider two different cell strains, Cell_1_ and Cell_2_, in mixed culture. Cell_1_ produces signal molecule *S*_1_ and Cell_2_ produces signal molecule *S*_2_. In each cell, toxin *T* is produced by the activation of signal molecules released by cells of its own type. The antitoxin *A* is actively produced by signal molecules released by cells of the other type. The antitoxin *A* sequesters the toxin *T* and forms a stable complex *TA*\* to repress the death process.

We set the relative population between Cell_1_ and Cell_2_ to unity 1 for demonstration. When cell strain Cell_1_ has a larger population than Cell_2_, more *S*_1_ than *S*_2_ will be synthesized and released into the environment. Signal molecules *S*_1_ will then diffuse into cells of both strains. In Cell_1_, toxin *T* will be produced in higher amount than antitoxin *A*, so the population of Cell_1_ will decrease. The opposite occurs in Cell_2_ since there is a higher amount of antitoxin *A* than toxin *T*. This stops cells in strain Cell_2_ from dying. As a result, the population of Cell_1_ decreases and the population of Cell_2_ increases until they are equal. This feedback control loop using two orthogonal quorum sensing systems ensures mutual population tracking and enforces the relative ratio between Cell_1_ and Cell_2_ to be one at steady state.

### D. The dual loop control strategy

The dual loop control strategy is illustrated in Figure 1c. The total population size and the relative ratio are independently set by two reference signals. It is necessary that the three quorum sensing molecules *S_g_, S*_1_, and *S*_2_ are mutually orthogonal to avoid crosstalk.

We introduce the feedback control for the two controller modules in the dual loop, which requires species that act to effectively annihilate or stabilize each other in biochemical reactions at either RNA or protein level. For example, *D* sequesters *G* and *A* sequesters *T* to form functionless complexes.

## III. THE GLOBAL REGULATION LOOP

### A. The biochemical reactions model

The deterministic model for global regulation corresponding to the biochemical reactions in Table I is derived according to mass-action and Michaelis-Menten kinetics. We make the following assumptions:

**TABLE 1.** Species, parameters and biochemical reactions in the global regulation

| Species | Description |  |
| --- | --- | --- |
| $C$ | Cell population | |
| $G$ | Species deciding the rate of cell growth and division | |
| $D$ | Species strongly binding with and annihilating $G$ | |
| $S_g$ | Global quorum sensing signal molecules | |
| Parameters | Description | Values |
| $k_C$ | Cell growth rate constant | $0.01 \mu\text{M}^{-1}\text{min}^{-1}$ |
| $C_{max}$ | Carrying capacity for cell growth | $10^5 \text{ mL}^{-1}$ |
| $\gamma_C$ | Cell dilution rate constant | $0.01 \text{ min}^{-1}$ |
| $K_S$ | Dissociation constant for $S_g$ | $2 \mu\text{M}$ |
| $\beta$ | Hill function coefficient | 2 |
| $k^+$ | Binding rate of effective annihilation | $0.02 \mu\text{M}^{-1}\text{min}^{-1}$ |
| $g_G$ | Basal production rate of $G$ | $0.3 \mu\text{Mmin}^{-1}$ |
| $g_D$ | Basal production rate of $D$ | $0.01 \mu\text{Mmin}^{-1}$ |
| $k_D$ | Maximal production rate of $D$ | $0.2 \mu\text{Mmin}^{-1}$ |
| $d_G$ | Dilution rate of $G$ | $0.01 \text{ min}^{-1}$ |
| $d_D$ | Dilution rate of $D$ | $0.01 \text{ min}^{-1}$ |
| $c_S$ | Synthesis rate of $S_g$ | $0.0002 \mu\text{MmLmin}^{-1}$ |
| $d_S$ | Degradation rate of $S_g$ | $0.25 \text{ min}^{-1}$ |
| $\gamma_S$ | Dilution rate of $S_g$ | $0.01 \text{ min}^{-1}$ |
| Reactions | Description |  |
| $\emptyset \xrightarrow{k_C GC(1-\frac{C}{c_{\max}})} C$ | Logistic growth of cell | |
| $\emptyset \xrightarrow{g_G} G$ | Production of species $G$ by basal induction | |
| $\emptyset \xrightarrow{g_D+k_D\text{Hill}(S_g)} D$ | Production of species $D$ by basal induction and activation by $S_g$ | |
| $G + D \xrightarrow{k^+} GD^*$ | Annihilating binding of $G$ and $D$ | |
| $G \xrightarrow{d_G} \emptyset, \quad D \xrightarrow{d_D} \emptyset, \quad S_g \xrightarrow{d_{S_g}} \emptyset$ | Dilution of $G, D$ and degradation of $S_g$ | |
| $C \xrightarrow{\gamma_C} \emptyset, \quad S_g \xrightarrow{\gamma_{S_g}} \emptyset$ | Dilution of $C$ and $S_g$ in chemostat | |

- Every cell in the population contains an identical negative feedback loop.
- Cell growth follows logistic kinetics with growth rate constant *k_C_* and carrying capacity *C*_max_, and the growth rate is proportional to the concentration of the growth regulating species *G*.
- There is dilution of the cell population and signal molecules, because the entire experiment is assumed to take place in a chemostat.
- The production of a species *x* is characterized by its basal and maximal rates *g_x_, k_x_*.
- Activation by regulator *x* is governed by a Hill function with dissociation constant *K*_*x*_ and Hill coefficient *β*_*x*_.
- Effective annihilation is achieved under the assumption that the binding reaction is much faster than the unbinding reaction and the complex is difficult to degrade.
- All species are assumed to decay with first-order kinetics.
- The synthesis of signal molecules *S_g_* occurs at a constant rate and *S_g_* reaches quasi-steady state by fast diffusion and degradation. Fast degradation can be implemented enzymatically as in [25].

We obtain the following model:

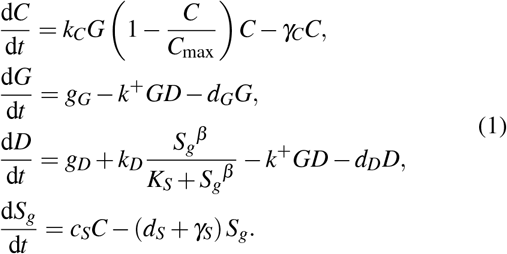

### B. The lag compensator

Let *C*_0_ be the total population reference. It is set by tuning the basal induction rates *g_G_* and *g_D_* of *G* and *D*, according to the equation:

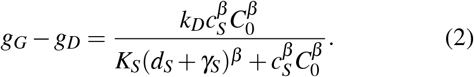

We remark that the conditions 0 < *C*_0_ < *C*_max_ and *g_D_* < *g_G_* < *g_D_* + *k_D_* must hold for the reference *C*_0_ to exist. In other words, it is not possible to tune the feedback controller to an arbitrary reference signal [26].

Assuming a feasible reference signal *C*_0_, let *S*_*g*_0__ and *S_g_* be the corresponding quasi-steady states of the signal molecules. Then we can define the tracking error *e_glo_* in global regulation as

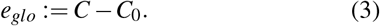

The quasi-steady states of the signal molecules are then derived as

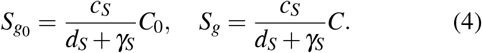

Thus, we obtain that

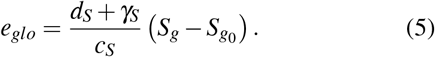

To emphasize the input term in our controller, we define

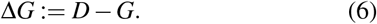

By subtracting the corresponding equations that describe the dynamics of *G* and *D* in equation (1), we can obtain that

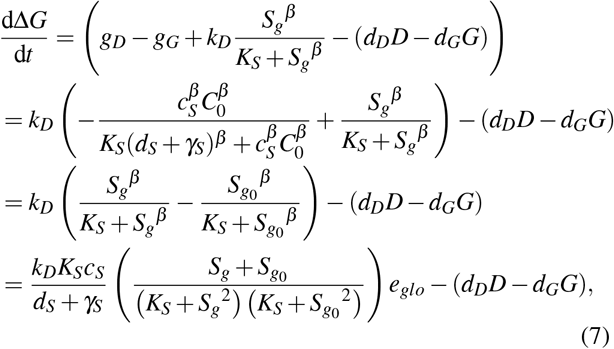
where for simplicity, we have assumed *β* = 2.

Then equations (1) and (7) set up the following dynamical system:

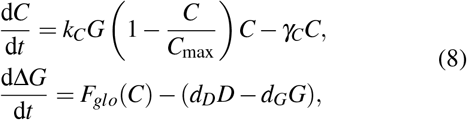
where 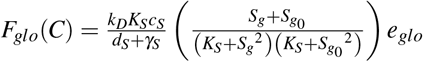.

In order to achieve perfect adaptation and be an integral controller, the control input dynamics should only be a function of the state *C* [26]. However, as Ang and McMillen note, this is not realistic for biological systems when protein degradation and dilution are present. Here, the rates *d_D_* and *d_G_* encompass both the degradation and the dilution processes. While we may assume that the degradation of *G* and *D* takes place at a low rate and can be approximated to 0, their dilution rate must equal the cell growth rate 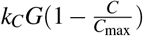. Thus, equation (8) is equivalent to

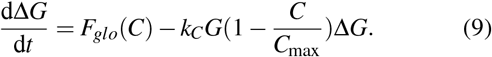

The feedback implemented in our system will be a lag compensator. It can be tuned to become closer to integral control by decreasing the cell growth rate *k_C_G*. When cells divide slowly, the error will decrease. For a cell division time of 60 minutes, *k_C_G* ≈ 0.01 min^−1^. We remark that the controller will have *e_glo_* ≈ 0 at steady state given 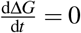.

### C. Local stability of the global regulation loop

We convert the ODE model of global regulation into a linearized state space model to better understand the proposed lag compensator and to examine the local stability of the closed-loop system. We assume that the regulated cell population is much smaller than carrying capacity, so the cell population is only regulated by the proposed controller, i.e. *C* ≪ *C*_max_. Also, the dilution rates of *G* and *D* are *d* = *d_G_* = *d_D_* = *k_C_G*, so the linearized state space model is in the following form:

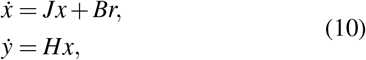
where the state *x*, input *r*, output *y*, Jacobian matrix *J*, input matrix *B* and output matrix *H* are given by

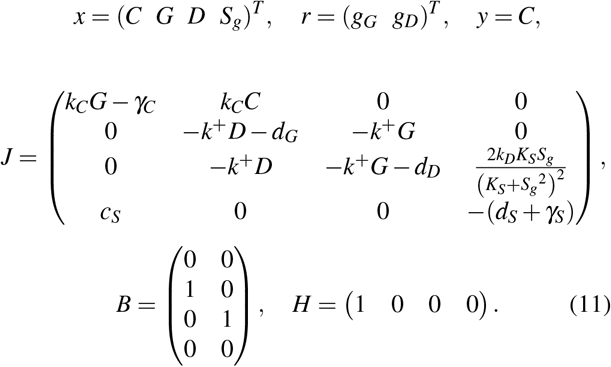

To determine the local stability of the global regulation, we can derive eigenvalues from the Jacobian matrix. They are computed as −0.076, −0.120, −0.065 + 0.065*j*, −0.065 −0.065*j*, which all lie in the left half space. Hence, the global regulation loop is locally stable.

### D. The tracking function performance ofthe controller

To demonstrate that the global regulation loop maintains the total population of cells, we simulate the dynamics of total population *C* with different induction rates *g_G_* of *G*, as illustrated in Figure 2a. Furthermore, we perturb the cell growth rate of one of the strains at time *t* = 1500 min and show robustness and adaptation in the closed loop system. We compare this with the performance of the open loop system in Figure 2b. Here, the steady state is only bounded by the carrying capacity of the consortium. The global regulation demonstrates set-point tracking of the reference as well as adaptation to a perturbation in the cell growth rate.

**Fig. 2.**
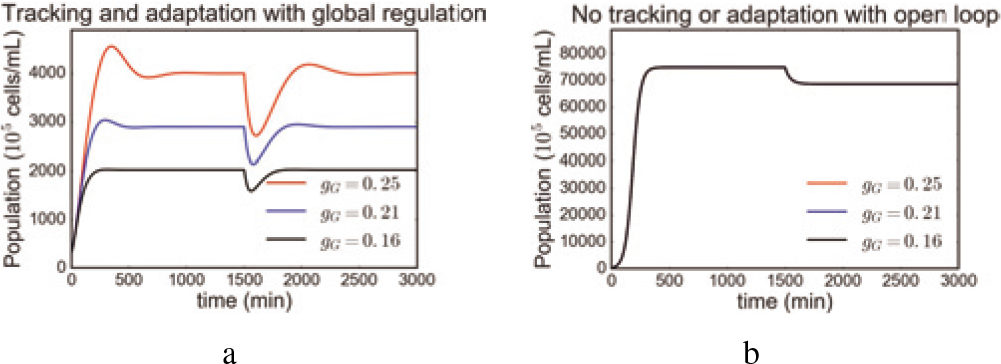
The simulated cell population dynamics with global regulation control show set-point tracking and adaptation. a. The set-point tracking function with global regulation responds to reference signals set by tuning the induction rate *g_G_*. After perturbing the growth rate of one cell strain, the total population recovers to the set-point value, which shows that global regulation enables robustness and adaptation. b. No tracking or adaptation occurs in the open loop system.

## IV. THE CO-REGULATION LOOP

### A. The biochemical reaction model

We consider cell strains Cell1 and Cell2 in mixed culture. Species, parameters and biochemical reactions are listed in Table II. We list the additional assumptions of the co-regulation model:

- There is an identical negative feedback loop in individual cells of the same strain, Cell1 or Cell2.
- The cell death rate is proportional to the concentration of the toxin *T* in the cell with constant *d_c_*.
- The quorum sensing systems are orthogonal.

**TABLE II.** Species, Parameters and Biochemical Reactions in Co-Regulation

| Species | Description |  |
| --- | --- | --- |
| $C_i$ | Cell population of strain $i$ | |
| $T^{(i)}$ | Toxin in cell strain $i$ | |
| $A^{(i)}$ | Antitoxin in cell strain $i$ | |
| $S_i$ | Quorum sensing signal molecules $i$ | |
| Parameters | Description | Values |
| $k_{C_i}$ | Cell growth rate constant for both strains | $0.04 \text{ min}^{-1}$ |
| $d_C$ | Cell death rate constant | $0.05 \text{ mM}^{-1} \text{ min}^{-1}$ |
| $g_T$ | Basal production rate of $T$ | $0.28 \text{ } \mu\text{Mmin}^{-1}$ |
| $k_T$ | Maximal production rate of $T$ | $0.2 \text{ } \mu\text{Mmin}^{-1}$ |
| $g_A$ | Basal production rate of $A$ | $0.05 \text{ } \mu\text{Mmin}^{-1}$ |
| $k_A$ | Maximal production rate of $A$ | $0.2 \text{ } \mu\text{Mmin}^{-1}$ |
| $d_T$ | Dilution rate of $T$ | $0.01 \text{ min}^{-1}$ |
| $d_A$ | Dilution rate of $A$ | $0.01 \text{ min}^{-1}$ |
| Reactions | Description |  |
| $\varnothing \xrightarrow{k_{C_i} C_i \left(1 - \frac{C_1 + C_2}{C_{\max}}\right)} C_i$ | Logistic growth of cell strain $i$ | |
| $C_i \xrightarrow{d_C T^{(i)}} \varnothing$ | Death of cell strain $i$ by toxin killing | |
| $\varnothing \xrightarrow{g_T + k_T \text{Hill}(S_i)} T^{(i)}$ | Production of toxin $T^{(i)}$ by basal induction and activation by $S_i$ | |
| $\varnothing \xrightarrow{g_A + k_A \text{Hill}(S_j)} A^{(i)}$ | Production of antitoxin $A^{(i)}$ by basal induction and activation by $S_j$ | |
| $T^{(i)} + A^{(i)} \xrightarrow{k^+} TA^{*(i)}$ | Annihilating binding of $T^{(i)}$ and $A^{(i)}$ | |
| $T^{(i)} \xrightarrow{d_T} \varnothing$,<br>$A^{(i)} \xrightarrow{d_A} \varnothing$,<br>$S_i \xrightarrow{d_S} \varnothing$ | Dilution of $T^{(i)}$, $A^{(i)}$ and degradation of $S_i$ | |
| $C_i \xrightarrow{\gamma_C} \varnothing$, $S_i \xrightarrow{\gamma_S} \varnothing$ | Dilution of $C_i$ and $S_i$ in chemostat | |

We derive the model for {*i, j*} = {1,2} as

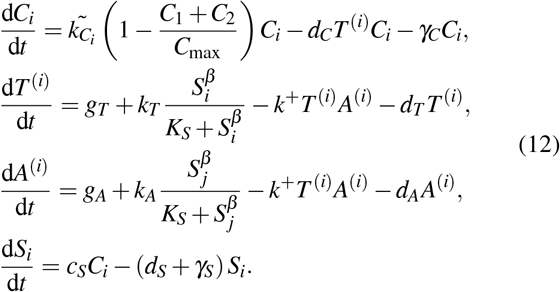

### B. The lag compensator

We first remark that the toxin and antitoxin species *A*^(*i*)^ and *T*^(*i*)^ dilute because of cell division. Hence, 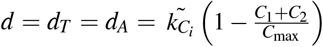. The lag compensator occurs when strong sequestration happens between *T* and *A* in the toxin-antitoxin system.

Let the relative population ratio between Cell_1_ and Cell_2_ be set to one, which defines a mutual tracking function. Then we can define the tracking error in the co-regulation as

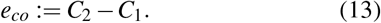

Consider *S*_1_ and *S*_2_ be the corresponding quasi-steady states of the signal molecules and they are then derived as

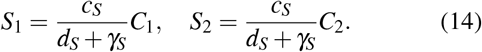

Thus, we obtain that

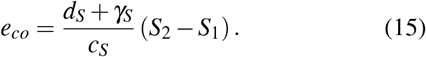

To emphasize the controller in Cell_2_, we define

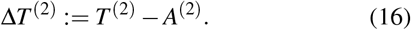

By subtracting the corresponding equations describing dynamics of *T*^(2)^ and *A*^(2)^ in equation (13), we can obtain

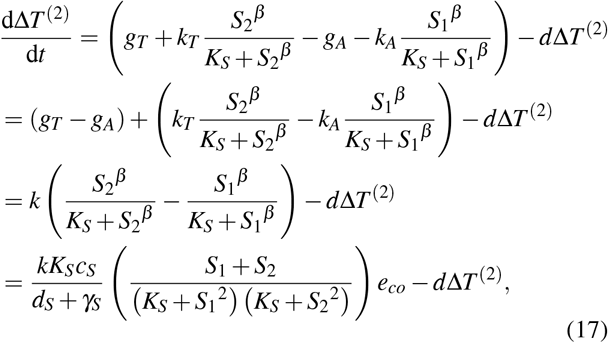
where we set *g_A_* = *g_T_* = *g, k_A_* = *k_T_* = *k* for the relative population ratio to be one and *β* = 2.

Equations (12) and (17) set up the dynamical system for Cell_2_:

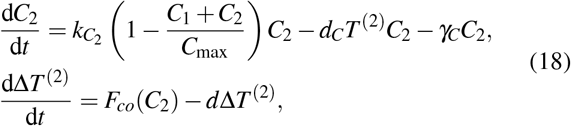
where 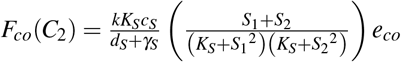.

The feedback implemented in co-regulation is also a lag compensator, and when cells divide slowly, we will have *e*_*co*_ = *C*_2_ − *C*_1_ ≈ 0 at steady state given 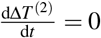.

### C. Cell population dynamics depend on their growth and death rates

The dynamics of the cell populations depend on the cell growth rate *k_C_* and the death rate *d_C_*. The steady states of the two cell populations can switch to oscillations from fixed stable points when growth rate increases or death rate decreases. The simulation results of cell populations *C*_1_ and *C*_2_ illustrated in Figures 3a and 3b show that cells must grow slowly to prevent oscillations. The Hopf bifurcation diagrams demonstrate that the dynamics switch when *k*_*C*_ > 0.07 *μ*M^−1^min^−1^ and *d*_*C*_ < 0.025 *μ*M^−1^min^−1^, as in Figures 3c and 3d.

**Fig. 3.**
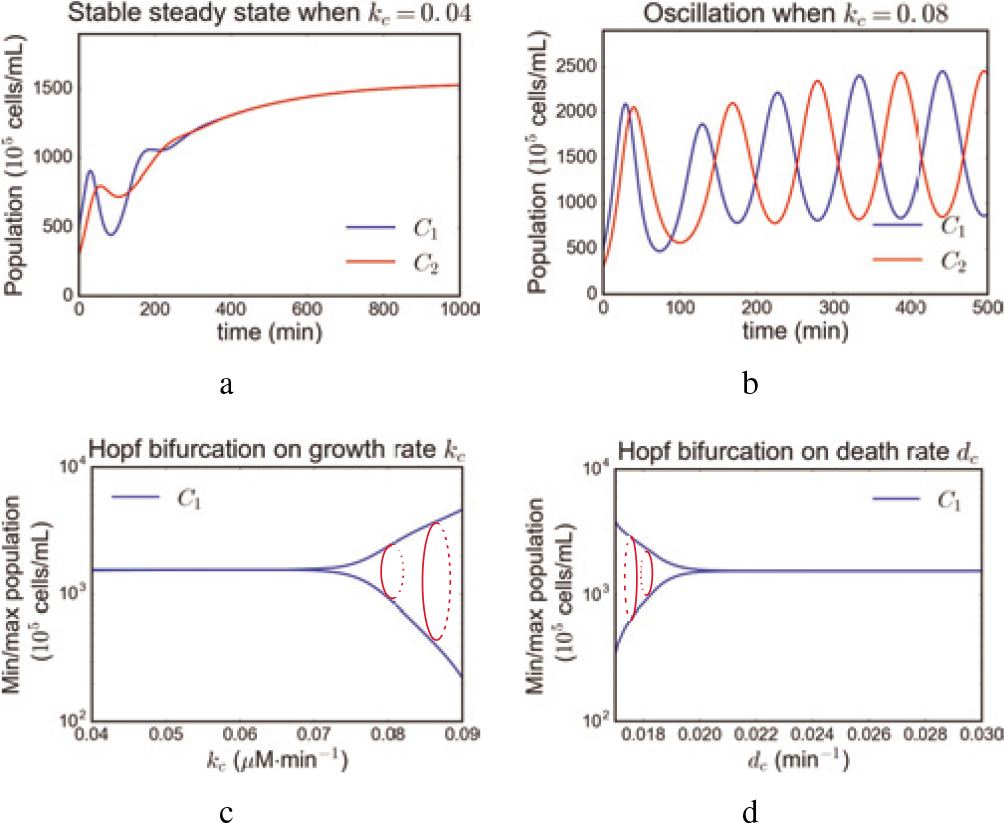
Cell must grow slowly to prevent oscillations. a. The dynamics of cell populations Cell_1_ and Cell_2_ converge to a stable steady state with relative ratio one when the growth rate *k_C_* = 0.04 *μ* M^−1^ min^−1^. b. The dynamics of cell populations Cell_1_ and Cell_2_ are oscillatory when the growth rate *k_C_* = 0.08 *μ*M^−1^min^−1^. c. The Hopf bifurcation diagram of the growth rate *k_C_* illustrates that when *k_C_* increases, the dynamics switch to oscillations from a fixed stable point. d. The Hopf bifurcation diagram of the death rate *d_C_* illustrates that when *d_C_* decreases, the dynamics switch to oscillations from a fixed stable point.

This oscillatory behavior in this two-strain mutual tracking system is caused by the delay one cell strain experiences in following population changes in the other one. When one cell strain grows fast because its growth rate is high or its death rate is low, there is a longer relative delay before the quorum sensing module is fully settled and regulates the production of toxin or antitoxin for tracking current state. This results in oscillations in both strains. Since the tracking is mutual, the oscillation of cell strains Cell_1_ and Cell_2_ is of same frequency and of a 180 degree phase difference.

## V. THE DUAL CONTROL LOOP

### A. The controller performance and robustness

To assess the behavior of the dual control loop system in response to internal set-point references on the total population and the relative population ratio, we define performance metrics of stability, response sensitivity, robustness, and adaptation to disturbances.

The set-point reference of the total population *C*_*tot*_ is fixed. We start with an initial condition (*C*_1_ (0), *C*_2_ (0), *C*_*tot*_ (0)) at time *t*_0_ = 0. We only consider scenarios when cell population converges to steady state after a time period *T*. At time *t*_1_ in the interval [*t*_0_, *t*_0_ + *T*], we add perturbations on the growth rate or we change the cell numbers of one cell strain and we observe the resulting dynamics of population. The metrics are defined as

- steady state error 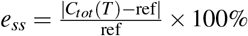,
- rise time *t*_rise_: the first time when *C*_*tot*_ (*t*_rise_) = ref,
- overshoot 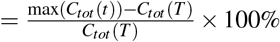
- recovery time *t*_rec_: time after t1 when it is true that 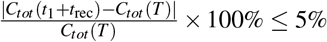.

In simulation, we set the total population to 3000 × 10^5^ cells/mL and the relative ratio of the two cell strains to value 1. At time *t*_1_ = 1500 min, we perturb the cell growth rate or, alternatively, we perturb the cell numbers. The consortium recovers to the previous steady state after an extrinsic perturbation on the absolute cell numbers. When we introduce a perturbation of 20% of the growth rate *k*_*C*_1__ of Cell1 and measure the performance metrics, we illustrate in equation (19) that the steady state error is almost zero for the total population. Thus, the lag compensator of the global regulation fulfills its function. Meanwhile, the co-regulation shows mutual population tracking between the two cell strains and maintains their relative ratio at value 1 in steady state.

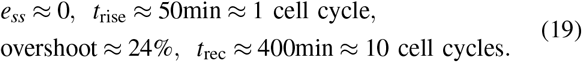

Furthermore, we vary the amplitude of the perturbation on rate *k*_*C*_1__ from 0% to 100%, we measure the population steady states of cell strains Cell_1_, Cell_2_, and of the total population *C_tot_*. Figure 4 shows that the steady state of total population always recovers to 3000 × 10^5^ cells/mL and the relative ratio also returns to value 1 for perturbations of less than 80%. This demonstrates that the dual loop regulation is robust and adapts to perturbations on the cell growth rate.

**Fig. 4.**
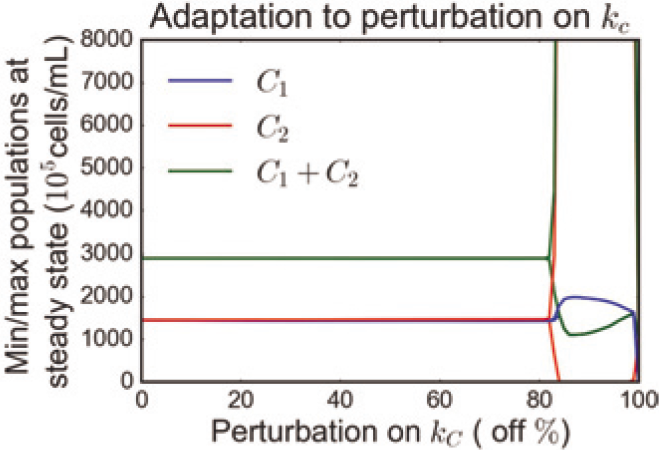
Perturbations on the growth rate of one cell strain. The dual loop regulation controller shows adaptation to perturbations on cell growth rate *k*_*C*_1__ of Cell_1_. The steady state of the total population and of the individual populations are maintained at 3000 × 10^5^ cells/mL and 1500 × 10^5^ cells/mL, respectively. When the growth rate perturbation is larger than 80%, the dual regulation controller fails.

### B. Independent tuning of the total population and of the relative cell strain ratio

We can choose total population and relative ratio reference signals and verify that the dual loop control strategy ensures separate tuning to the two reference signals. Figure 5 shows that both regulation functions are realized with small steady state error at representative total population and relative ratio reference values.

**Fig. 5.**
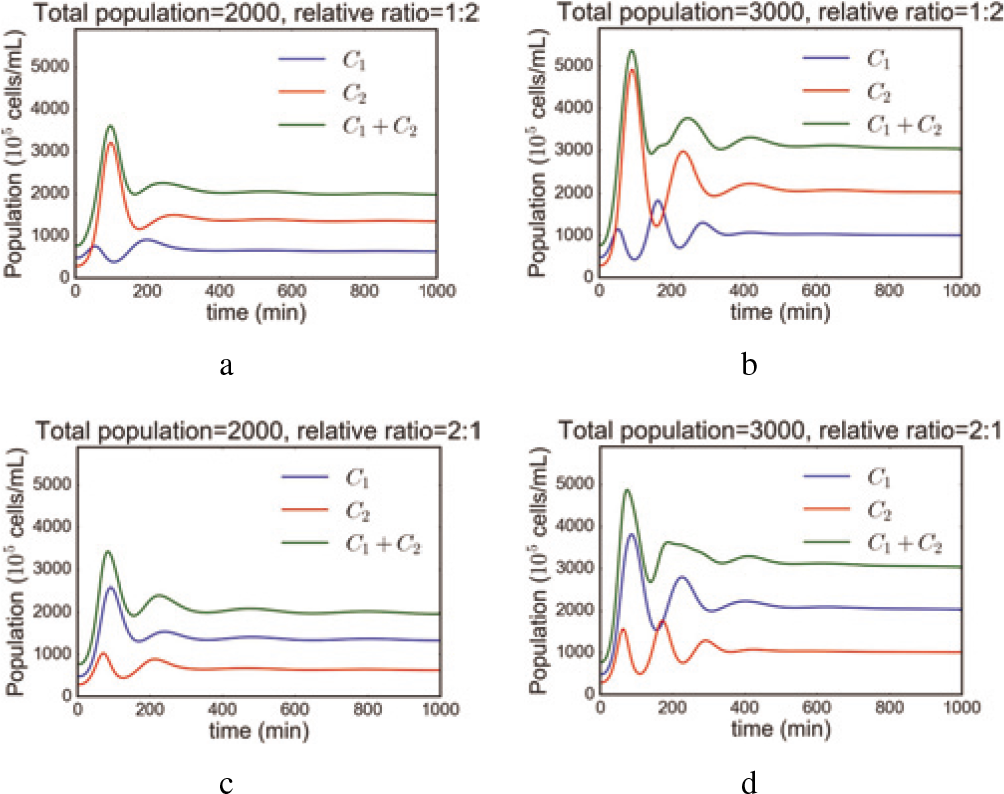
The dual loop controller dynamics with independently tuned values of the total population and of the relative ratio of the cell strains. a. Reference signals *C*_1_ + *C*_2_ = 2000, *C*_1_/*C*_2_ = 1: 2. b. Reference signals *C*_1_ + *C*_2_ = 3000, *C*_1_/*C*_2_ = 1: 2. c. Reference signals *C*_1_ + *C*_2_ = 2000, *C*_1_/*C*_2_ = 2: 1. d. Reference signals *C*_1_ + *C*_2_ = 3000, *C*_1_/*C*_2_ = 2:1.

## VI. DISCUSSION

In this paper, we considered a dual lag compensator to separately regulate the total population size and the relative population ratio of two cell strains in a microbial consortium. The general control strategy of dual loop controller design can be applied to any synthetic systems with sensors, comparators, and actuators. We proposed a mathematical model for the dual loop regulation by considering reactions and parameters from the synthetic biology literature and we implemented the resulting circuit in *in-silico* experiments. Our simulation results demonstrate that dual lag compensator control enables set-point tracking with adaptation for a range of total population and relative ratio reference signals. We investigated the robustness of the closed-loop system by assessing its adaptation to perturbations on the cell growth rate and also changes in cell number. The introduced response metrics of the system are representative of a realistic environment.

We provide design guidelines and predict experimental results in microbial consortia. We are constructing corresponding biological circuits and measuring preliminary data based on synthetic tools such as the ccdB/ccdA toxin-antitoxin system, gp2/RNA antisense technology and AHL quorum sensing pathways in *E. coli*. In our future work, we will include the stochasticity in biochemical processes in our models and controller designs and we will carry out experimental biological implementation.

## VII. Acknowledgments

The authors would like to thank Reed McCardell and Victoria Hsiao for their insightful discussion. The authors X. R., A. B., and A. S. are partially supported by the Air Force Office of Scientific Research, grant number FA9550-14-1-0060. The project depicted is also sponsored by the Defense Advanced Research Projects Agency (Agreement HR0011-17-2-0008). The content of the information does not necessarily reflect the position or the policy of the Government, and no official endorsement should be inferred.

